# Host EPAC1 modulates rickettsial adhesion to vascular endothelial cells via regulation of ANXA2 Y23 phosphorylation

**DOI:** 10.1101/2021.04.20.440643

**Authors:** Zhengchen Su, Thomas Shelite, Yuan Qiu, Qing Chang, Maki Wakamiya, Jiani Bei, Xi He, Changcheng Zhou, Yakun Liu, Angelo Gaitas, Tais Saito, Bin Gong

**Author notes:** **Correspondence to:** Bin Gong, MD, PhD, or Tais B Saito, DVM, PhD.

## Abstract

Recently we have identified that endothelial surface annexin A2 (ANAX2) functions as a receptor for spotted fever group rickettsial adhesin outer membrane protein B (OmpB), which binds to the endothelial cell (EC) surface. Moreover, we reported that intracellular cAMP receptor EPAC1 modulates ANXA2 tyrosine (Y) 23 phosphorylation, and inactivation of EPAC1 suppresses ANXA2 expression on the EC luminal surface by downregulating Y23 phosphorylation. Since we reported that EPAC1 plays a critical role in the initial step to successfully establish rickettsial infection of ECs, this work aims to answer the following: (a) What is the mechanism underlying the regulatory role of EPAC1 in ECs during the initial step of bacterial infection? (b) Is the EPAC1-ANXA2 signaling pathway involved in the regulation of rickettsial adhesion to ECs?

In the present study, an established system that is anatomically-based and quantifies bacterial adhesion to ECs *in vivo* was combined with novel fluidic force microscopy (FluidFM) to dissect the functional role of the EPAC1-ANXA2 signaling pathway in rickettsiae–EC adhesion. We reveal that the deletion of the *EPAC1* gene impedes rickettsial binding to endothelium *in vivo*. In addition, single living brain microvascular EC study that employs FluidFM and site-directed mutagenesis provides evidence that supports our finding that EPAC1 governs rickettsial adhesion to EC surfaces via regulation of ANXA2 Y23 phosphorylation.

## Introduction

The mechanism(s) underlying bacterial adherence to vascular endothelial cells (ECs) under shear stress from flowing blood is critical to our understanding the initial stages of the pathogenesis of endovascular bacterial infections. The constitutively expressed native proteins on and/or in the mammalian host plasma membrane play crucial functional role(s) during the initial stage of infection, just prior to EC activation. Recently we reported that endothelial surface annexin A2 (ANAX2) functions as a receptor for spotted fever group (SFG) rickettsial adhesin outer membrane protein B (OmpB) binding and is also involved in establishing *Staphylococcus aureus* adhesion to EC surfaces (*1*). Moreover, we reported that intracellular cAMP receptor <u>e</u>xchange <u>p</u>rotein directly <u>a</u>ctivated by <u>c</u>AMP 1 (EPAC1) modulates ANXA2 tyrosine (Y) 23 phosphorylation, and inactivation of EPAC1 suppresses ANXA2 expression on the EC luminal surface by downregulating Y23 phosphorylation (*2*). Since our discovery that EPAC1 plays a critical role in the initial step of a successfully established rickettsial infection of ECs (*3*), we are now extending this work to answer the following questions: (a) What is the mechanism underlying the regulatory role of EPAC1 in ECs during the initial step of bacterial infection? (b) Is the EPAC1-ANXA2 signaling pathway involved in the regulation of rickettsial adhesion to ECs?

In the present study, an established, anatomically based, *in vivo* quantitative system that measures bacterial adhesion to ECs (*1*) was combined with novel fluidic force microscopy (FluidFM) to dissect the functional role of the EPAC1-ANXA2 signaling pathway during initial adhesion of rickettsiae to EC surfaces. We reveal that the deletion of the *EPAC1* gene impedes rickettsial binding to endothelium *in vivo*. A study coupling FluidFM and site-directed mutagenesis with a single, living brain microvascular EC (BMEC) provides evidence that supports our finding that EPAC1 governs rickettsial adhesion to EC surfaces via regulation of ANXA2 Y23 phosphorylation.

## Results

### 1. *In vivo* corroborating rickettsial adhesion to EC surfaces in an EPAC1-dependent manner

Our previous work provided *in vitro* and *ex vivo* evidence that EPAC1 plays a critical regulatory role in the early stage of rickettsial invasion into nonphagocytic host cells (*3*). Using our recently established *in vivo* quantitative system to measure bacterial adhesion to vascular Ecs (*1*) in our *EPAC1*-knock out (KO) mouse model (*2*), we can now verify the regulatory role of EPAC1 during rickettsial adhesion. Plaque assays reveal that the number of viable rickettsiae in circulating blood was higher in *EPAC1*-KO mice (n=11) than wild-type (WT) mice (n=10) at 30 min and 1 hr post-infection (p.i.) when the bacteria were given by the intravenous route (**Fig. 1A**). Similar as our previous report (*1*), immunofluorescence staining showed that, in WT mice, rickettsiae were mainly detected on the luminal surface or tunica intima of the vascular wall (arrows in **Fig. 1B**). However, in *EPAC1*-KO mice, fewer rickettsiae were detected in the same areas. In contrast, unattached rickettsiae were visible in blood clots in the lumen of these blood vessels (arrow heads in **Fig. 1B**).

**Figure 1:**
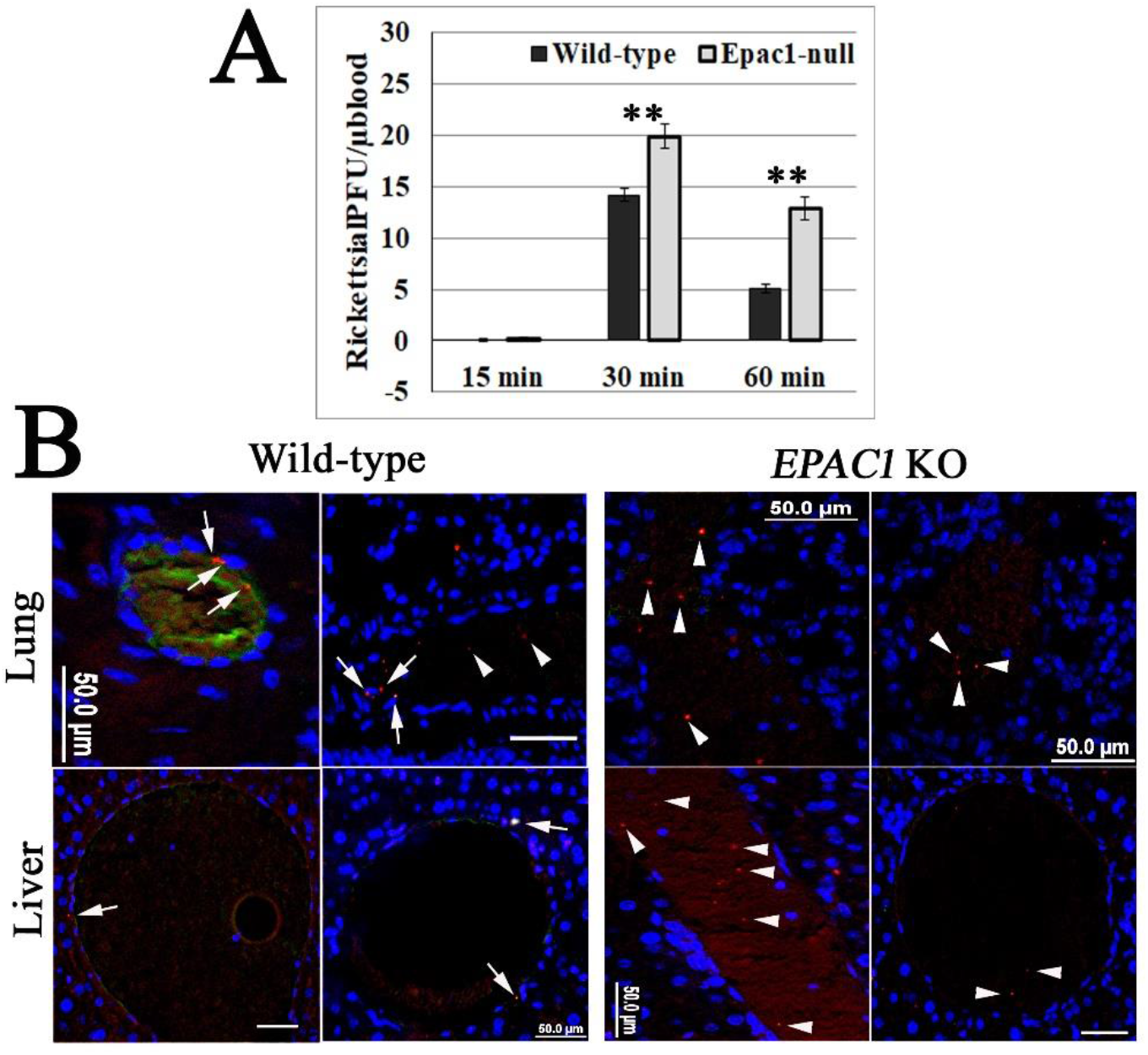
Global depletion of *EPAC1* blocks rickettsial adherence to blood vessel luminal surfaces *in vivo*. **A**: Plaque assay for *R. australis* using blood samples collected from the orbital venous sinus of wild-type (WT) (n=10) and *ANXA2*-knock out (KO) (n=11) mice at different times p.i. with 10 LD_50_ doses of *R. australis* given by the intravenous route. Data are represented as mean ± SEM. **: compared to the control group, P < 0.01. **B**: Representative IF-based identification of rickettsiae (red) in WT and *EPAC1*-KO mice 60 min p.i. with 10 LD_50_ doses of *R. australis* given by the intravenous route. EPAC1 is stained with green signals. Rickettsiae adhere to the intima layer of blood vessels (arrows) or wrap in blood clots (arrowheads) in the lumens of blood vessels. Nuclei are stained blue. Scale bars, 50µm.

These observations suggest that rickettsial adhesion to the vascular luminal surface occurs in a host EPAC1-dependent manner.

### 2. Rickettsial OmpB shows a host EPAC1-dependent binding strength on the surface of a living EC

Employing our *in vivo* model and atomic force microscopy (AFM), we found that rickettsiae mediate adhesion to the surface of ECs via ANXA2 (*1*). Compared to traditional AFM using a protein-functionalized colloid cantilever to evaluate the protein–protein interaction, the micropipette cantilever of FluidFM enables us to use exchangeable colloid probes with the same cantilever for the quantification of irreversible interacting forces, drastically increasing the experiment throughput. Moreover, it reduces the bias from excessive usage of a single functionalized colloid (**Fig. 2**) (*4*).

**Figure 2:**
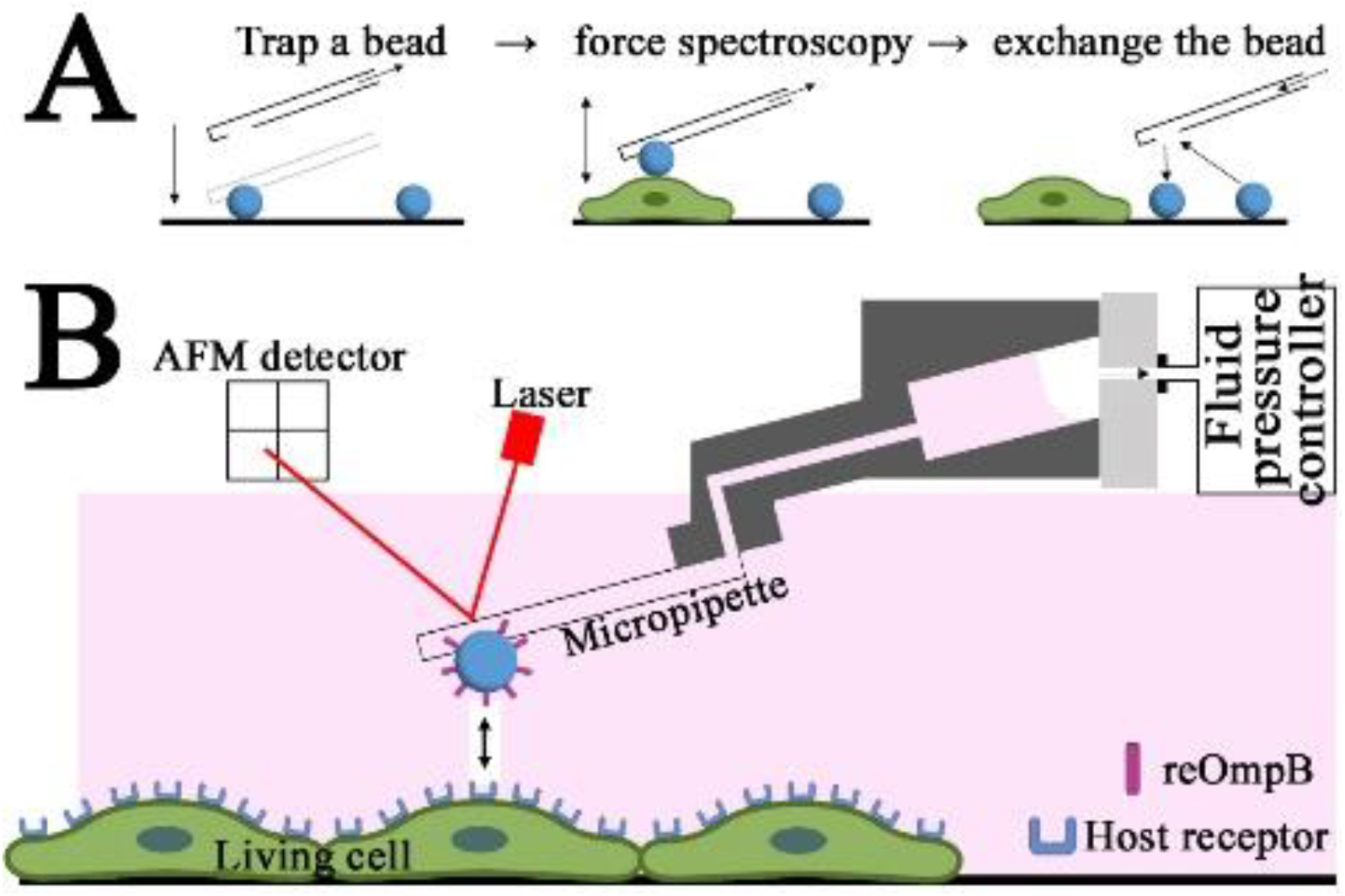
Fluidic force microscopy (FluidFM). (A) Operation moods of using exchangeable colloid probes with the same cantilever for the quantification of irreversible interacting forces. (B) Schematic representation of FluidFM using micropipette-based cantilever.

OmpB is a rickettsial adhesin responsible for host invasion (*5, 6*). Using recombinant OmpB (reOmpB)-functionalized microbeads, we employed the force spectroscopy capacity of the FluidFM system to determine the binding strength between reOmpB and the surface of single living BMEC (**Fig. 3**). BSA coated microbeads were used as negative control. As expected, reOmpB-coated microbeads generally exhibit higher adhesion forces to WT BMECs (P < 0.01). To examine whether EPAC1 contributes to reOmp and BMEC binding, we compared the adhesion forces generated in WT versus *EPAC1*-KO BMECs; reduced adhesion forces between a reOmpB-coated microbead and a *EPAC1*-KO BMEC were recorded (P < 0.01) (**Fig. 3**).

**Figure 3:**
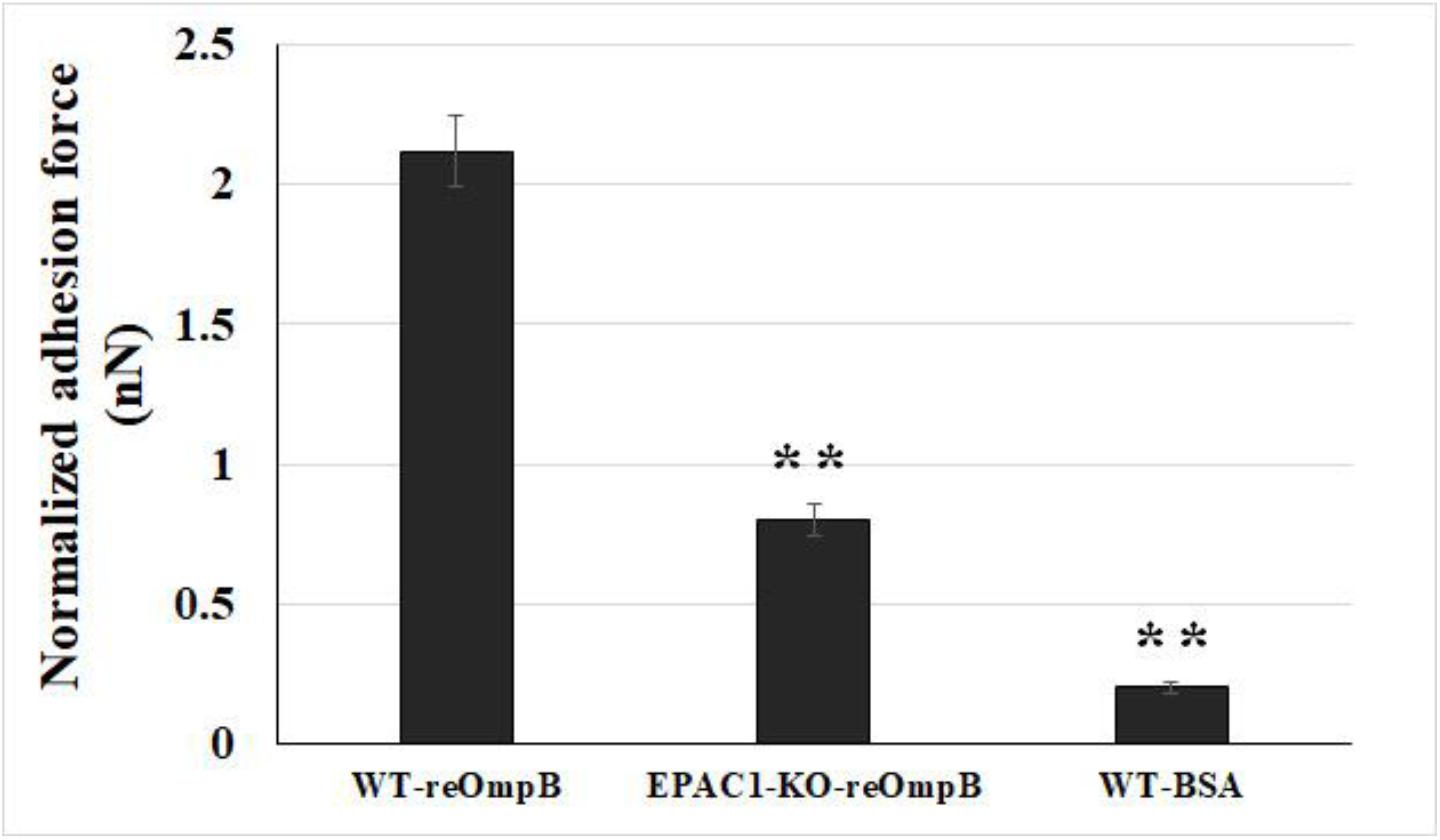
Rickettsial OmpB shows a host EPAC1-dependent binding strength on the surface of a living EC. FluidFM studies measure the binding forces (nanoNewton, nN) between reOmpB-coated microbead and a single living WT or *EPAC1*-KO mouse BMEC. Data are represented as mean ± SEM. **: compared to the group of WT, P < 0.01. At least three different detection areas were measured in one cell. Ten cells were sampled per group.

Furthermore, taking advantage of a recently developed non-cAMP analogue, an EPAC1 specific activator named I942 (*7, 8*), we observed that I942 enhanced the interacting force between reOmpB and a live WT BMEC (P<0.05) (**Fig. 4**). Depletion of *ANXA2* in ECs diminished reOmpB binding forces to the *ANXA2*-KO BMEC surface (P<0.01) (**Fig. 4**), corroborating our previous report that anti-ANXA2 antibody weakens the interacting force between reOmpB and a living EC (*1*). Interestingly, depletion of *ANXA2* in ECs reduced the enhanced binding force between reOmpB and ECs in the presence of I942 (versus the WT group, p=0.09).

**Figure 4:**
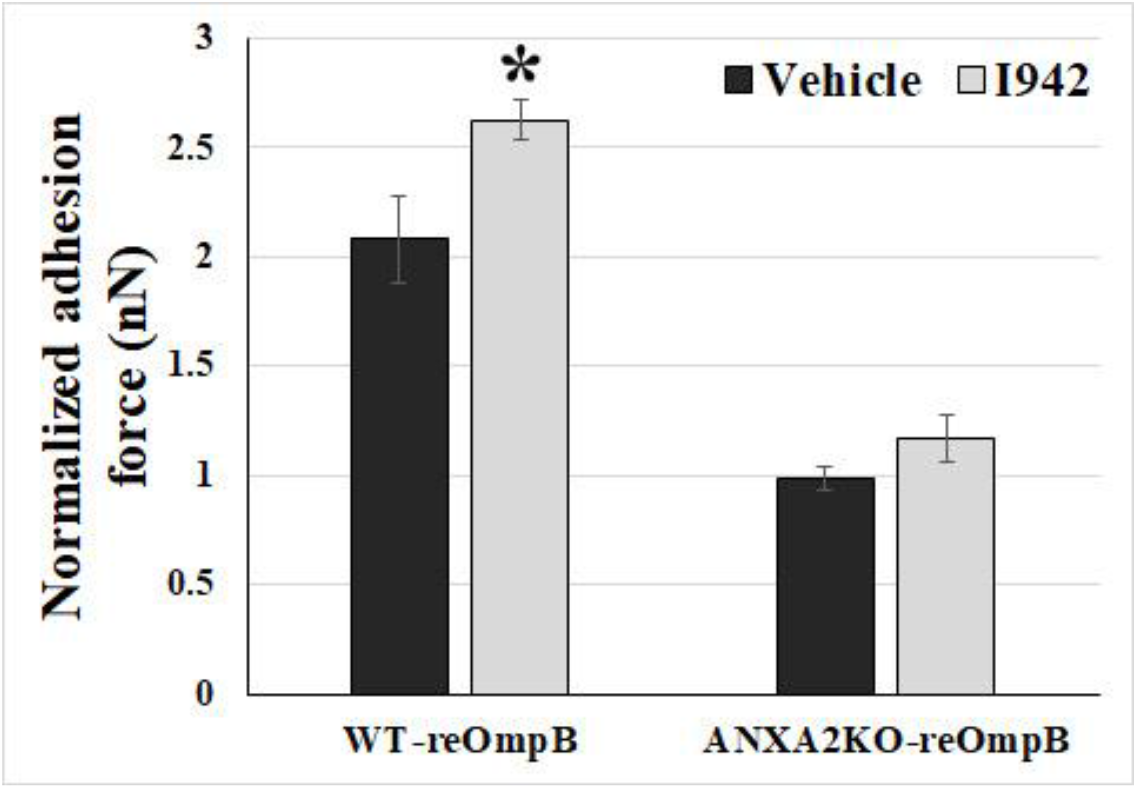
EPAC1 specific activator enhances reOmpB binding force to single living EC dependent of ANXA2. FluidFM studies measured the binding forces (nanoNewton, nN) between reOmpB-coated microbead and single living WT or *ANXA2*-KO mouse BMEC, which was exposed to I942 (5uM) or vehicle for 24 hr. Data are represented as mean ± SEM. *: compared to the group of vehicle, P < 0.05. At least three different detection areas were measured in one cell. Ten cells were sampled per group.

Collectively, these data suggest that the EPAC1-ANXA2 pathway is involved in the interaction of OmpB with ECs during the initial stages of bacterial binding.

### 3. Ectopic expression of phosphodefective and phosphomimic mutants replacing Y23 of ANXA2 in *ANXA2*-KO BMECs results in the display of different binding force to reOmpB

Our previous study suggested that EPAC1 regulates endothelial luminal surface ANXA2 expression by modulating the phosphorylation of Y23 (*2*). To identify whether Y23 is an EPAC1-targeted site in the N-terminus of ANXA2 that leads to regulation of rickettsial adhesion, we transfected *ANXA2*-KO BMECs with mouse WT *ANXA2* construct, phosphodefective *ANXA2* mutant Y23F, or phosphomimic *ANXA2* mutant Y23E, respectively, which were generated by site-directed mutagenesis using their respective mutant oligonucleotides (*9*). Ectopic expression of the WT construct (versus no construct group, p=5.36 × 10^−8^) and phosphomimic Y23E (versus no construct group, p=6.70 × 10^−5^) in *ANXA2*-KO BMECs rescued the diminished reOmpB and cell binding force, respectively, whereas the phosphodefective mutant Y23F (p=0.90) failed to do so (**Fig. 5**). Furthermore, ectopic expression of the mouse WT *ANXA2* construct also rescued EPAC1-specific activator I942-induced enhancement of the binding force in *ANXA2*-KO BMECs, while no such effect in the empty vector controls.

**Figure 5:**
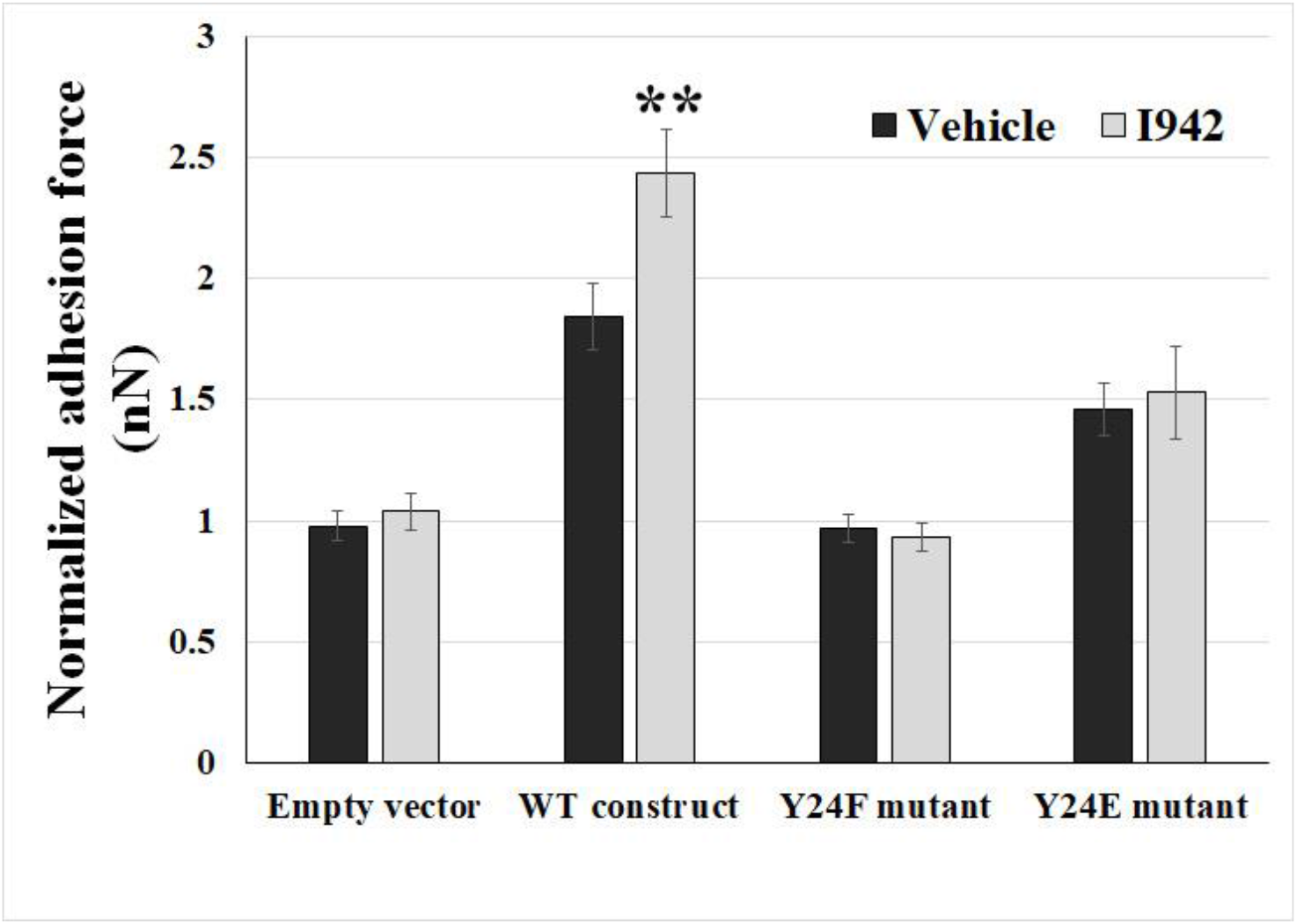
Host EPAC1 governs rickettsial adhesion to EC surface via regulation on phosphorylation of Y23 in ANXA2. FluidFM studies measured the binding forces (nanoNewton, nN) between reOmpB-coated microbead and single living *ANXA2*-KO mouse BMEC, which was transfected with WT *ANXA2* construct, phosphodefective *ANXA2* mutant Y23F, or phosphomimic *ANXA2* mutant Y23E, after exposure to I942 (5uM) or vehicle for 24 hr. The cells were transfected with empty vectors were used as negative controls. Data are represented as mean ± SEM. **: compared to the group of vehicle, P < 0.01. At least three different detection areas were measured in one cell. Ten cells were sampled per group.

Collectively, these data suggest that host EPAC1 governs rickettsial adhesion to EC surfaces via regulation of phosphorylation of Y23 in ANXA2.

## Discussion

As an obligate intracellular bacterium, rickettsia is required to initiate adherence to host cells prior to infection. OmpB is the most abundant membranous protein in SFG rickettsia and functions as an adhesin during attachment and invasion (*10*). Recently, we reported that EPAC1 plays a critical role during rickettsial invasion (*3*) and identified endothelial ANXA2 as a novel receptor for rickettsial OmpB binding to (*1*). In the present study, we corroborated rickettsial adhesion to EC surfaces in an EPAC1-dependent manner *in vivo*. Our FluidFM assay, employing an exchangeable colloid-based micropipette, provided further evidence of host EPAC1-dependent binding of rickettsial OmpB on the surface of a living EC. In addition, our single living cell study that coupled FluidFM and site-directed mutagenesis showed that Y23 in the N-terminus of ANXA2 is an EPAC1-targeted site involved in modulating rickettsial adhesion.

ANXA2 in the cellular membrane compartment is believed to be regulated mainly via phosphorylation/dephosphorylation of the N-terminal domain of ANXA2, including S11, Y23, and S25 (*11, 12*). Phosphorylation of Y23 was identified as a regulatory switch during association of ANXA2 with lipid rafts (*2, 13*) and translocation of ANXA2 between different subcellular compartments (*2, 14*). After phosphorylation of Y23 via the Src family tyrosine kinase(s)-dependent pathway, ANXA2 is translocated to the cell surface in the format of heterotetramer with S100A10 by a yet unknown mechanism (*13-15*). We reported that inactivation of EPAC1 down-regulates Y23 phosphorylation of ANXA2, associated with ANXA2 endothelial surface translocation, and does not impact the constitutive level of vWF in the plasma or other major hematological parameters (*2*). Specifically, another ANXA2 phosphodefective mutant Y23A, not ANXA2 phosphomimic mutant Y23E, yields a predominantly negative effect for the translocation of ANXA2 to the membrane surface. The Y23A mutant can still bind to S100A10 but only in the cytosol, not in the plasma membrane (*16*). Given that ANXA2 functions as a binding receptor for rickettsial OmpB, in the present study we further reveal that EPAC1 regulates rickettsial adhesion to EC surfaces by targeting the ANXA2 Y23. Depletion of *ANXA2* in ECs reduced the enhanced binding force between reOmpB and ECs in the presence of I942. The WT *ANXA2* construct rescued EPAC1-specific activator I942-induced enhancement of reOmpB binding forces to a ANXA2-KO BMEC surface. Furthermore, Y23E mutant rescued the diminished reOmpB and cell binding force in *ANXA2*-KO BMEC, whereas the Y23F mutant failed.

The colloidal probe-based AFM assay was developed to study protein–protein interactions and to quantify the interacting forces in the range from picoNewtons to nanoNewtons (*4*). During manufacture, a colloid probe of different sizes is attached to a tipless cantilever by gluing and coated with recombinant proteins or antibodies by the user (*1, 2*). Entire colloidal probe-based cantilevers must be exchanged between experiments to measure different biomolecular interactions on the same sample with different protein functionalization. Contamination and degradation of colloid surfaces limits the lifetime of a colloid probe and induces potential bias when the probe is reused to measure the same biomolecule in a different sample. FluidFM was developed to conquer these limitations. A microfluidic pressure controller applies negative pressure to a connected hollow cantilever to quickly grab a functionalized microbead, and the microbead can be released by applying positive pressure through the micropipette channel (*4*). Thus, FluidFM offers the following advantages: (1) it allows for the use of exchangeable colloids for the quantification of irreversible and long-term protein–protein interactions, (2) it provides strong connections between the colloid and cantilever, (3) rapid exchange of microbeads with different protein coatings is possible, and (4) the method consistently uses the same reflex side of a single cantilever with different colloidal probes (*4*). In the present study we use a reOmpB-coated microbead-based FluidFM cantilever to quantify the interaction between reOmpB and single, living cell in multiple experimental groups.

## Conclusion

We have revealed a novel mechanism by which EPAC1 modulates rickettsial adhesion, involving regulation of tyrosine 23 phosphorylation of the binding receptor ANXA2. This finding provides experimental and theoretical support for therapeutic strategies for rickettsiosis targeting the host cAMP-EPAC system.

## Materials and Methods

### Mice

All animal experiments were performed according to protocols approved by the Institutional Animal Care and Use Committee of the University of Texas Medical Branch (UTMB). Wild-type (WT) mice (C57BL/6J) were obtained from The Jackson Laboratory (Bar Harbor, ME). C57BL/6 EPAC1-KO mice were derived as described previously (*2*). *ANXA2*-KO mice on the C57BL/6J background were a generous gift from Dr. Katherine Hajjar (Weill Cornell Medicine, NY, NY) (*17*). All mice used in this study were 8- to 12-week-old males. C57BL/6J mice are highly susceptible to *R. australis*. Therefore, this organism was chosen as the SFG rickettsial agent of choice (*18*).

### *Rickettsia* and cell culture

*R. australis* (strain Cutlack) was prepared as described (*3*). Uninfected Vero cells were processed as mock control material using the same procedure. All biosafety level (BSL-) 3 or ABSL-3 experiments were performed in CDC-certified facilities in the Galveston National Laboratory at UTMB (Galveston, TX), using established procedures and precautions.

Mouse brain microvascular endothelial cells (BMECs) were isolated from wild-type, *EPAC1*-KO, and *ANXA2*-KO mice using established protocol(*1, 8, 19*).

### Antibodies and other reagents

A rabbit polyclonal antibody against SFG rickettsiae was obtained from the laboratory of Dr. David Walker (UTMB, Galveston, TX). AlexaFluor 594-conjugated goat anti-rabbit IgG and ProLong Gold Antifade Reagent with DAPI were purchased from Invitrogen (Carlsbad, CA). Recombinant rickettsial OmpB (reOmpB) (amino acids 1363-1655) was purchased from MyBioSource (San Diego, CA). Unless otherwise indicated, all reagents were purchased from Thermo Fisher Scientific. Expression constructs for GFP-tagged mouse WT ANXA2, ANXA2 mutant Y23E, and ANXA2 mutant Y23F were purchased from GENEWIZ (South Plainfield, NJ).

### Cell transfection

Transfections were performed using a published protocol (*11*). Cells were transfected with of WT ANXA2, ANXA2 Y23E, and ANXA2 Y23F constructs (1µg/500 µL for 1 × 10^5^ cells), respectively, using Magnetofection PolyMag & PolyMag Neo (OZ Biosciences, San Diego, CA). The efficacy of transfection was evaluated with GFP fluorescent microscopy (*11*).

WT construct sequence:

ATGTCTACTGTCCACGAAATCCTGTGCAAGCTCAGCCTGGAGGGTGATCATTCTACACCCCCAAGTGCCTACGGGTCAGTCAAACCCTACACCAACTTCGATGCTGAGAGGGATGCTCTGAACATTGAGACAGCAGTCAAGACCAAAGGAGTGGATGAGGTCACCATTGTCAACATCCTGACAAACCGCAGCAATGTGCAGAGGCAGGACATTGCCTTCGCCTATCAGAGAAGGACCAAAAAGGAGCTCCCGTCAGCGCTGAAGTCAGCCTTATCTGGCCACCTGGAGACGGTGATTTTGGGCCTATTGAAGACACCTGCCCAGTATGATGCTTCGGAACTAAAAGCTTCCATGAAGGGCCTGGGGACTGACGAGGACTCCCTCATTGAGATCATCTGCTCACGAACCAACCAGGAGCTGCAAGAGATCAACAGAGTGTACAAGGAAATGTACAAGACTGATCTGGAGAAGGACATCATCTCTGACACATCTGGAGACTTCCGAAAGCTGATGGTCGCCCTTGCAAAGGGCAGACGAGCAGAGGATGGCTCAGTTATTGACTACGAGCTGATTGACCAGGATGCCCGGGAGCTCTATGATGCCGGGGTGAAGAGGAAAGGAACCGACGTCCCCAAGTGGATCAGCATCATGACTGAGCGCAGTGTGTGCCACCTCCAGAAAGTGTTCGAAAGGTACAAGAGCTACAGCCCTTATGACATGCTGGAGAGCATCAAGAAAGAGGTCAAAGGGGACCTGGAGAACGCCTTCCTGAACCTGGTCCAGTGCATCCAGAACAAGCCCCTGTACTTCGCTGACCGGCTGTACGACTCCATGAAGGGCAAGGGGACTCGAGACAAGGTCCTGATTAGAATCATGGTCTCTCGCAGTGAAGTGGACATGCTGAAAATCAGATCTGAATTCAAGAGGAAATATGGCAAGTCCCTGTACTACTACATC

CAGCAAGACACCAAGGGTGACTACCAGAAGGCACTGCTGTACCTGTGTGGTGGGGATGACTGA.

For ANXA2 mutant Y23E, the sequence of bases encoding Y23 is replaced with GAA. For ANXA2 mutant Y23F, the sequence of bases encoding Y23 is replaced with UUC.

### Mouse *in vivo*, anatomically-based, quantitative rickettsial adhesion measurement system

The quantitative mouse *in vivo* rickettsial adhesion assay was performed as described (*1*). The principal concept of this anatomically-based inoculation model is that the more rickettsiae adhere to the luminal surfaces of the blood vessels, the fewer rickettsiae are detected in peripheral blood samples. Multiple visceral organs are fixed without perfusion rinse (to keep blood in the vessel lumens) for immunofluorescence (IF)-based histological studies to identify unattached rickettsiae, which are trapped in clots in vessel lumens (*1*). Briefly, after an ordinarily lethal dose of *R. australis* (1 × 10^7^ PFU/0.2µl; the LD_50_ is 1 × 10^6^ PFU) was injected through the tail vein, rickettsial virulence (by plaque assay) was measured in a 1 µl blood sample collected from the orbital venous sinus (OVS) at different times until 1 hr post-infection (p.i.). Rickettsial antigens were detected with a rabbit polyclonal antibody against SFG rickettsiae (1:1000) overnight at 4°C. Nuclei were counter-stained with DAPI. Fluorescent images were analyzed using an Olympus BX51 epifluorescence or Olympus IX81 confocal microscope.

### FluidFM adhesion assay

Carboxylate modified latex microbeads (5 µm CML Latex Beads, Invitrogen) were coated with reOmpB using published protocols (*20, 21*). Briefly, microbeads were suspended in 2-(N-morpholino) ethanesulfonic acid (MES) buffer. Freshly made, water-soluble 1-ethyl-3-(3-dimethylaminopropyl) carbodiimide (EDAC) in MES (50 mg/ml) was added to the beads and incubated for 30 minutes. The beads were then washed twice in MES buffer with centrifugation. EDAC-activated beads were incubated with purified reOmpB with gentle mixing at room temperature for 4 hr. The bead-protein mixture was centrifuged and the supernatant was removed. The beads were then washed with phosphate-buffered saline (PBS) three times to completely remove free protein. 0.4% glycine in 0.1M PBS was added to the beads for 10 min to deactivate the crosslinking, followed by one washing in 0.1M PBS. The beads were then resuspended in 0.1% glycine storage buffer. The coating procedure was performed under sterile conditions. Negative control BSA coated beads were created using the same procedure, but replacing reOmpB with BSA.

The FluidFM system coupling Nanosurf Core AFM (NanoSurf, Liestal, Switzerland) and Fluidic Pressure Controller (Cytosurge AG, Glattbrugg, Switzerland) was used for this assay. A micropipette (2um, 0.3N/m) (Cytosurge AG) was coated with 0.1 mg/ml poly(L-lysine)-*g*-poly(ethylene glycol) (PLL-g-PEG) to block nonspecific adhesions prior to calibration using the method of Sander *et al*. (*22*) and pre-set functions of the Nanosurf software. The air in the reservoir on the backside of the micropipette microchanel was removed by PBS prior to being connected to the Pneumatic Connector (Cytosurge AG). After subsequent connection to the Fluidic Pressure Controller, positive pressure (20 mBar) was applied to enable PBS to flow through the microchannel within the micropipette. The micropipette was then functionalized with an reOmpB-coated latex microbead by applying negative fluid pressure (−800 mBar) to trap the bead on the aperture of the micropipette. To measure the adhesion force between the layer of proteins that was coated on the surface of the bead and the surfae of the single living cell, the bead-loaded micropipette was driven to approach the cell monolayer with 2 nN as the setpoint force, and paused on the surface of the cell for 0.5 min. This defined time established the interaction on the surface of the living cell (*4*). The force spectroscopy was performed in the designated area (10 x 10 µm) to measure the unbinding force during rupture of the interaction between the ligands expressed in the designated areas at the apical surface of the single livning cell and the reOmpB-coated microbead (*2*). Ten cells were sampled per group. The bead was replaced following measurments of three cells to avoid nonspecific contamination.

## Statistical analysis

Values are reported as mean ± SEM. The data were analyzed using the Student’s *t*-test or One-Way ANOVA (Sigmaplot, Sigma Stat, Jandel Scientific Software, San Rafael, CA). P values are as follows: **P < 0.01 and *P < 0.05. Statistical significance is considered as P < 0.05.

## Acknowledgements

We gratefully acknowledge Dr. Edward Nelson for his contributions for establishing the capacity of the FluidFM system. We gratefully acknowledge Dr. Volker Gerke for providing human ANXA2 constructs for the preliminary study. We gratefully acknowledge Drs. David Walker, Juan Olano, and Jia Zhou for providing reagents and Dr. Donald Bouyer for BSL-3 facility support. We thank Dr. Kimberly Schuenke for her critical review and editing of the manuscript. This work was supported by NIH grant R01AI121012 (BG), R21AI137785 (BG), R21AI154211(BG), R03AI142406 (BG), and R21AI144328 (TS and BG). The sponsors had no role in the study design, data collection and analysis, decision to publish, or preparation of the manuscript.

## Authorship Contributions

BG and TS designed the study, performed experiments, analyzed data, and wrote the manuscript. ZS, TS, YQ, QC, MW, XH, and YL performed experiments. ZS, YQ, BJ, and CZ prepared experiments and analyzed data, and helped prepare the manuscript. AG designed the study and analyzed data.

